# BioVR: a platform for virtual reality assisted biological data integration and visualization

**DOI:** 10.1101/307769

**Authors:** Jimmy F. Zhang, Alex R. Paciorkowski, Paul A. Craig, Feng Cui

## Abstract

**Background**: The effective visualization and presentation of biological data is of critical importance to research scientists. The increasing rate at which experiments generate data has exacerbated the visualization problem. While biological datasets continue to increase in size and complexity, the shift to adopt new user interface (UI) paradigms has historically lagged. Consequently, a major bottleneck for analysis of next-generation sequencing data is the continued use of the UIs primarily inspired from single-type and small-scale data in 1990’s.

**Results**: We have developed BioVR, an easy-to-use interactive, virtual reality (VR)-assisted platform for visualizing DNA/RNA sequences and protein structures using Unity3D and the C# programming language. It utilizes the cutting-edge Oculus Rift, and Leap Motion hand detection, resulting in intuitive navigation and exploration of various types of biological data. Using *Gria2* and its associated gene product as an example, we present this proof-of-concept software to integrate protein sequence and structure information. For any residue of interest in the *Gria2* sequence, it can be quickly visualized in its protein structure within VR.

**Conclusions**: Using innovative 3D techniques, we provide a VR-based platform for visualization of DNA/RNA sequences and protein structures in aggregate, which can be extended to view omics data.

## Background

Most current methods such as the UCSC Genome Browser [1], Ensembl [2], and the Integrative Genomics Viewer [3] adopt 2D graph representations for visualizing biomolecular sequences. The advent of massively parallel sequencing (MPS) technologies in the past decade has revolutionized the field of genomics, enabling fast and cost-effective generation of a large amount of sequence data [4]. This technological innovation leads to the accumulation of vast quantities of data across various-omics fields, posing a tremendous challenge to scientists for effective mining of data to explain a phenomenon of interest. Virtual reality (VR) presents a unique opportunity to address the challenge of visualizing and manipulating complex biological datasets across multiple domains.

VR refers to a 3D simulated environment generated by a computer into which users are immersed, as opposed to a 3D rendering of a 2D display [5]. In such an environment, users can observe internal complexity of the data, gain better understanding about the relationship among different elements, and identify previously unappreciated links between them. For these reasons, VR has potential benefits in abstract information visualization, and many studies have shown that users tend to understand data better and more quickly in VR environments than in conventional 2D and 3D desktop visualization [6-8].

Modern VR is based on a low-cost, stereoscopic head-mounted display (HMD), such as the Oculus Rift, Google Cardboard and HTC Vive, which presents different images to each eye to achieve 3D perception of the scene. In the case of Oculus Rift, HMD is accompanied by a head-tracking system using an infrared camera, a 3-axis gyroscope, and a hand-tracking device such as the Leap Motion. These developments in VR are very attractive to use in biological visualization because they give users intuitive control of exploring and manipulating complex biological data.

Unsurprisingly, VR has been used for the visualization of biomolecular [9] and metabolic [10] networks, microscopy data [11], protein-ligand complexes [12], biological electron-transfer dynamics [13], whole genome synteny [14], and a whole cell [15]. Moreover, VR has numerous applications in medical education [16, 17] and research [18–23].

However, none of these tools deal with the fast-growing sequence and structural data of biomolecules. Here, we present a proof-of-concept application, BioVR, which is a VR-assisted platform to integrate and visualize DNA/RNA sequences and protein structures in aggregate. The *Rattus norvegicus Gria2* gene (Glutamate Ionotropic Receptor AMPA type subunit 2) was used as a test case because its DNA sequence (on Chromosome 2, NC_005101), mRNA sequence (NM_001083811) and protein structure (5L1B) are well defined. This work allows researchers to visualize the DNA/RNA sequences of *Gria2*, together with its protein structures in VR. It facilitates our understanding of the sequence-structure relationship of important residues of a protein. Our results suggest that the effective visualizations and user interfaces in VR help researchers quickly identify important genomic variants found through the MPS technologies, and represent a valid alternative to traditional web-based tools currently available for genomics, transcriptomics, and proteomics.

## Implementation

### Hardware Requirements

A VR-capable computer with the following specifications was used: (1) Intel Core i7-6700HQ Quad Core processor, (2) 6GB GDDR5 NVIDIA GeForce GTX 1060 graphics card, and (3) 8GB RAM. Oculus Rift with the following specifications was used: Oculus App (version 1.16.0.409144 (409268)) and device firmware (version 708/34a904e8da). To detect a user’s hands, Leap Motion Software (version: 3.2.0+45899), Leap Motion Controller (ID: LP22965185382), and Firmware (version: 1.7.0) were used.

### Development Tools

The VR-enabled desktop application was built using Unity3D (Unity), a game development environment with virtual reality capabilities (Supplementary Figure S1). It utilizes the C# programming language, which comes as part of Microsoft’s .Net software. Although both Unity and Unreal Engine support VR, Unity was chosen as the development platform for BioVR because it has ample online resources and is available on campus, through the virtual reality lab of the Center for Media, Arts, Games, Interaction and Creativity (MAGIC) of RIT.

A software package called UnityMol was created in Unity by Marc Baaden of Centre National de la Recherche Scientifique (CNRS). A non-VR 2014 version of this software, SweetUnityMol, is available for free download via SourceForge.net (Sweet UnityMol r676 beta r7) [24]. It is governed by a French copyright license called CeCILL-C which grants us the right to its free use and modification. With Unity version 5.4.2f, Sublime Text 3 Build 3126 (sublime), a code editor, and Git 2.11.0.windows.3 (git) for version control, we used the 2014 version (base code) as the basis of BioVR.

### Software Design and Development

#### (A) Basic Unity Objects

Software development within Unity constitutes its own specialty. A detailed discussion of Unity-specific challenges and best practices are beyond the scope of this paper. However, basic Unity concepts pertaining to BioVR are described below.

All objects in Unity are of type GameObject. A GameObject instance may contain zero or more components (Supplementary Table S1), including the MeshFilter. A MeshFilter object contains a reference to a Mesh instance. A Mesh instance can represent a single polyhedron by virtue of its internal data structures: vertices, triangles, normals, and UV coordinates (*i.e.*, coordinates scaled to the 0.1 range for texturing a image, with 0 representing the bottom/left of the image and 1 representing the top/right of the image) arranged in arrays of the appropriate type, which has an upper vertex limit around 6000. In BioVR, the primary objects of interest are GameObject instances which contain a Mesh instance. A mesh can be accessed via the following:

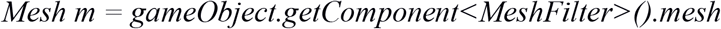

assuming there exists a non-null reference to GameObject named gameObject.

#### (B) Game Loop

The game loop is a ubiquitous concept in the gaming industry and influences BioVR in subtle but important ways. A game loop is a finite state machine that describes the high level game state at any given time point. It is modeled roughly as follows: when the player enters the game for the first time, the game loop starts. The current game state is rendered and displayed to the player. The player assesses the current game state and decides the next move, pressing the corresponding input(s). The next game state is computed based on the player’s inputs and scene information. As soon as the next game state is rendered on screen, it becomes the current game state. The player assesses the newly created game state and responds with inputs, repeating the loop. Unless game-ending conditions are triggered, the game loop continues.

Unity provides the MonoBehaviour class to help developers implement the game loop concept. The purpose of MonoBehaviour is to contain base code which organizes component actions into one of several game loop states. MonoBehaviour therefore contains the Awake(), Start(), and Update() functions, as well as other functions specific to Unity’s implementation of game loop design (Supplementary Table S2). To maintain game loop design principles, Unity encourages scripts to inherit from MonoBehaviour, but will otherwise compile and run normal C# classes with the caveat that those classes do not have the opportunity to directly affect game loop behavior.

VR applications rely on game loop architecture due to similarities in their state changes following user input. On start, the application takes on the state, *S_0_*, and information is rendered into the headset. Information about the player’s head orientation from the headset and finger positions from Leap Motion is gathered. The next state, *S_1_*, is then computed based on said information and rendered into the headset. Upon render completion, *S_1_* becomes the current game state, and input from the headset and Leap Motion determines the next state *S_2_*. This continues to the state *S_N_* so long as the user does not exit the application. The set of all states from *S_1_* to *S_N_* is part of the larger “playing” game state within the game loop. In BioVR, GameObjects that are subject to user input have some influence on the exact parameters of the next state and are therefore part of the game loop (Figure 1). These GameObjects have attached components that inherit from MonoBehaviour and define the Awake(), Start(), and Update() functions as appropriate.

**Figure 1.**
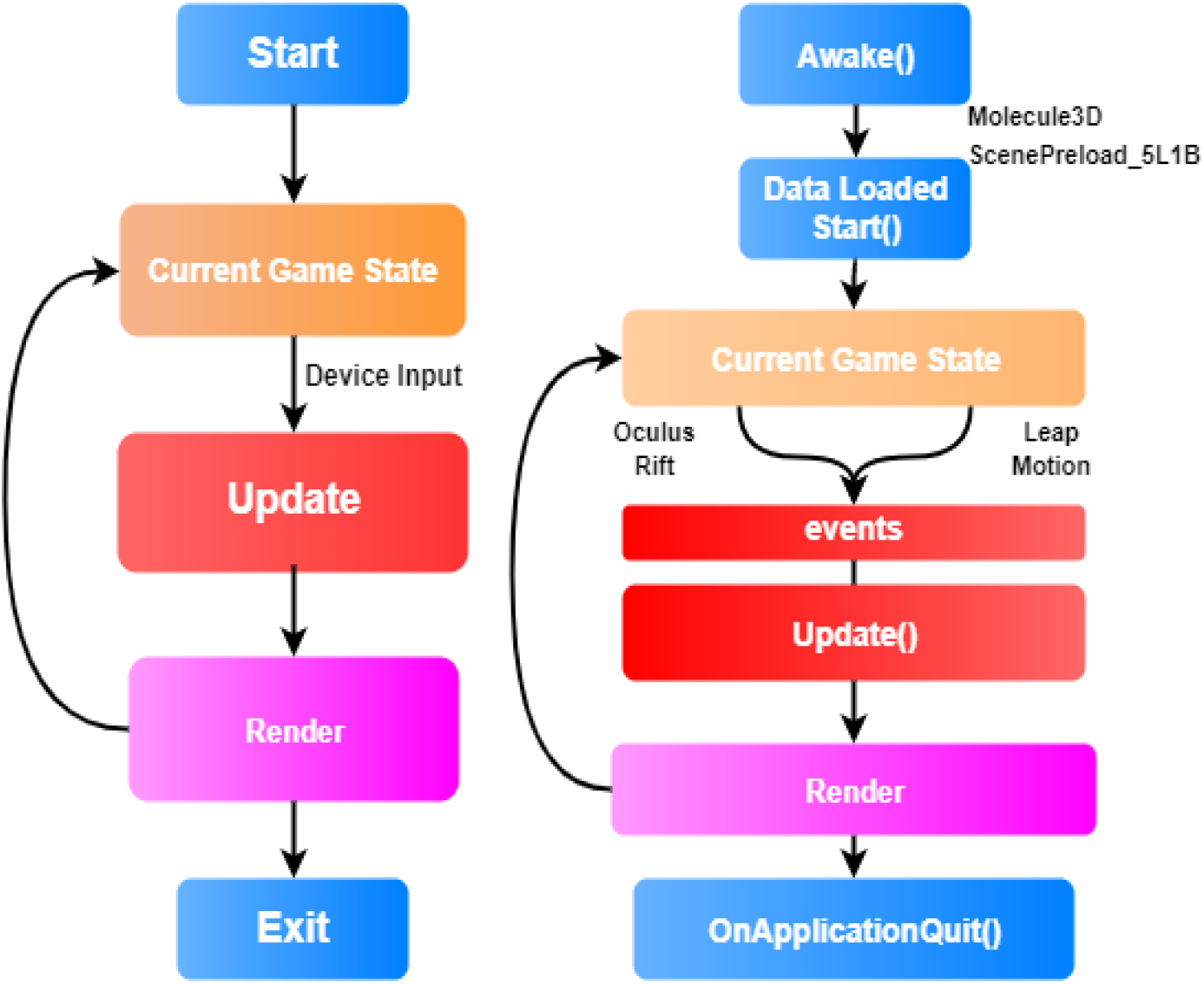
Comparison between the gaming operation and BioVR operation. (Left) A generic game loop. (Right) A high-level game loop specific to BioVR with important components that drive game state.

#### (C) MVC

Model-view-controller (MVC) design calls for the separation of concerns (SoC) into three different components of an application. The Model component is responsible for representing the data the application needs to interact with. The View component is primarily responsible for rendering and maintaining the correct views given inputs. The Controller mediates exchanges between Model and View components while also listening to user input events. The separation of concerns in MVC architecture is meant to keep application logic apart from their presentation. Changes and additions to base code rely heavily on the MVC design paradigm.

In the UnityMol (base code) package [24], modules primarily concerned molecular structure display reside within the*Molecule.View* folder, but can also be found in the *Molecule.SecondaryStructures* folder or within the surface folder without assignment to any particular namespace. Consequently, the MVC architecture in the base code resembles a Venn diagram (Figure 2) rather than cleanly separated MVC components (Supplementary Figure S2). In our subsequent software development, we did not attempt any code refactoring aimed at separating MVC components into their respective namespaces. Rather, separate namespaces were created as folders within “/Script/” apart from base code. Modules such as *Residue.cs, Nucleotide.cs, DNAPlaneController.cs*, were added to their respective namespaces.

**Figure 2.**
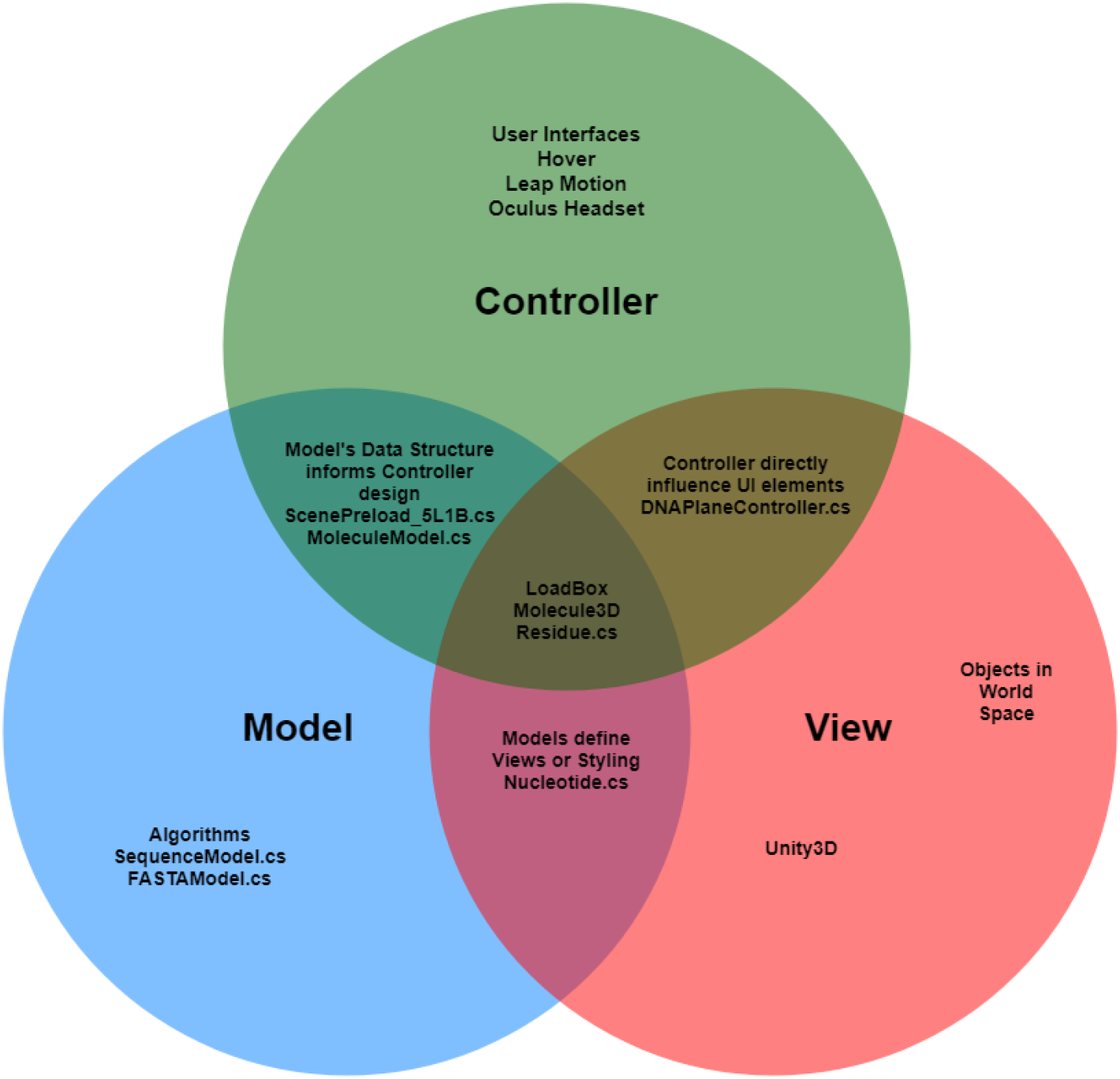
Major components of BioVR. Three separate namespaces including VRModel, View, and Controller were created to allow the Viewer faithfully execute on the MVC architecture. Note that certain components within the Viewer blur the line between different components of the ideal MVC model.

#### (D) Data Models & Inheritance

The use of inheritance throughout software development is a recommended best practice. It minimizes assumptions about common features of classes and their roles. In BioVR, object-oriented inheritance is utilized only to describe data models for parsing and holding biological data files (Figure 3).

**Figure 3.**
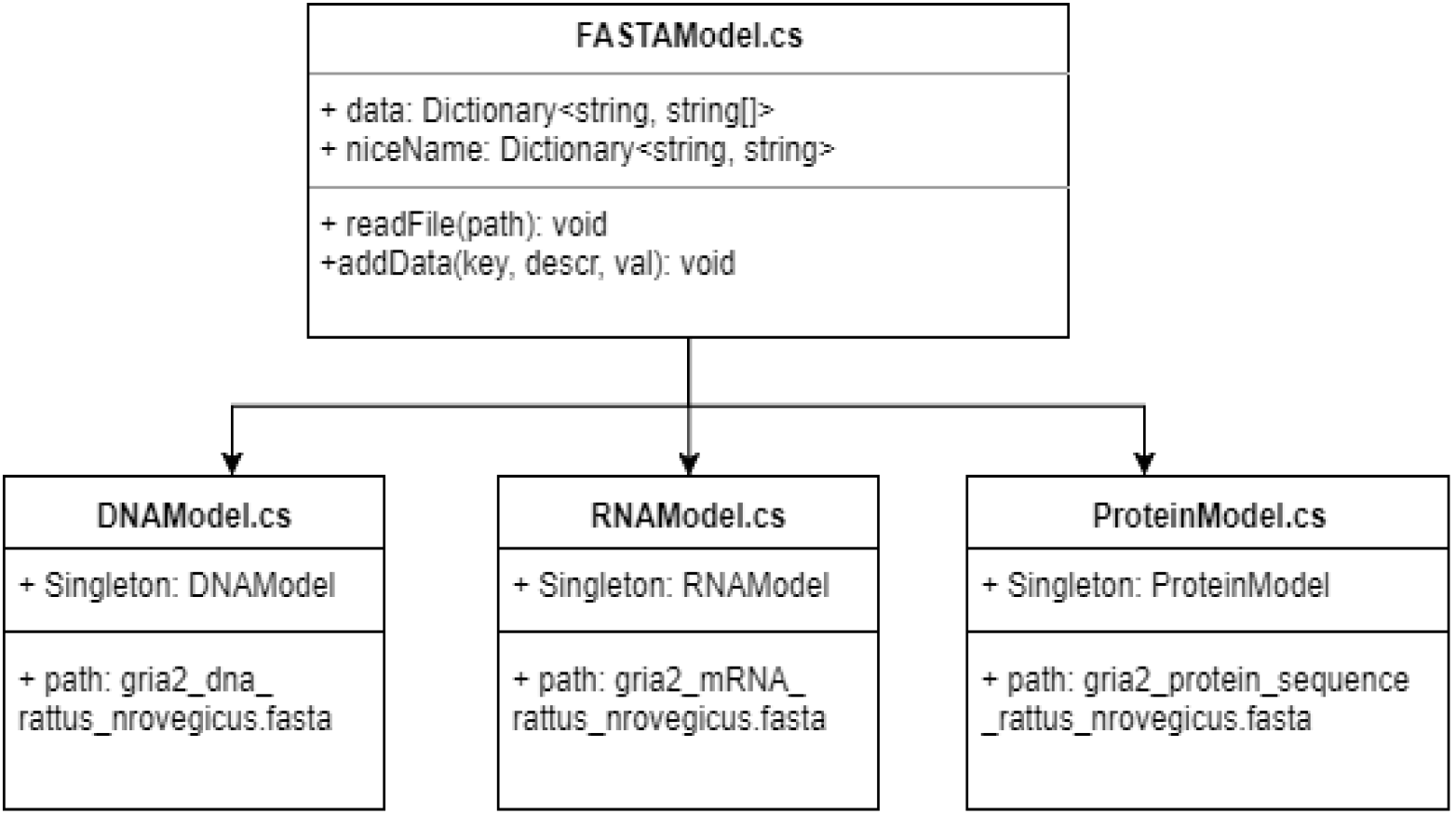
Inheritance relationship describing three different data models used in BioVR. The three data models, which are *DNAModel, RNAModel* and *ProteinModel,* are able to parse the data in the FASTA format. The *niceName* field maps common species names *(e.g. Homo sapiens)* to the key indices within the *data* field depending on the specific child class. The biological data of the *Gria2* gene (*e.g.,* DNA, mRNA and protein sequences) were used for the purpose of illustration.

All FASTA files are processed by the same way; therefore the common parsing code resides within the *FASTAModel.cs* module. Each species for which FASTA files are available map to a unique *niceName* string instance, e.g. string *niceName = “Rattus norvegicus”.* The *niceName* string instance is common to the DNA, RNA, and Protein models for each species, but they map to different valued keys within the data dictionary, depending upon which type of model (DNA, RNA, or Protein) that the user is interested (Figure 4).

**Figure 4.**
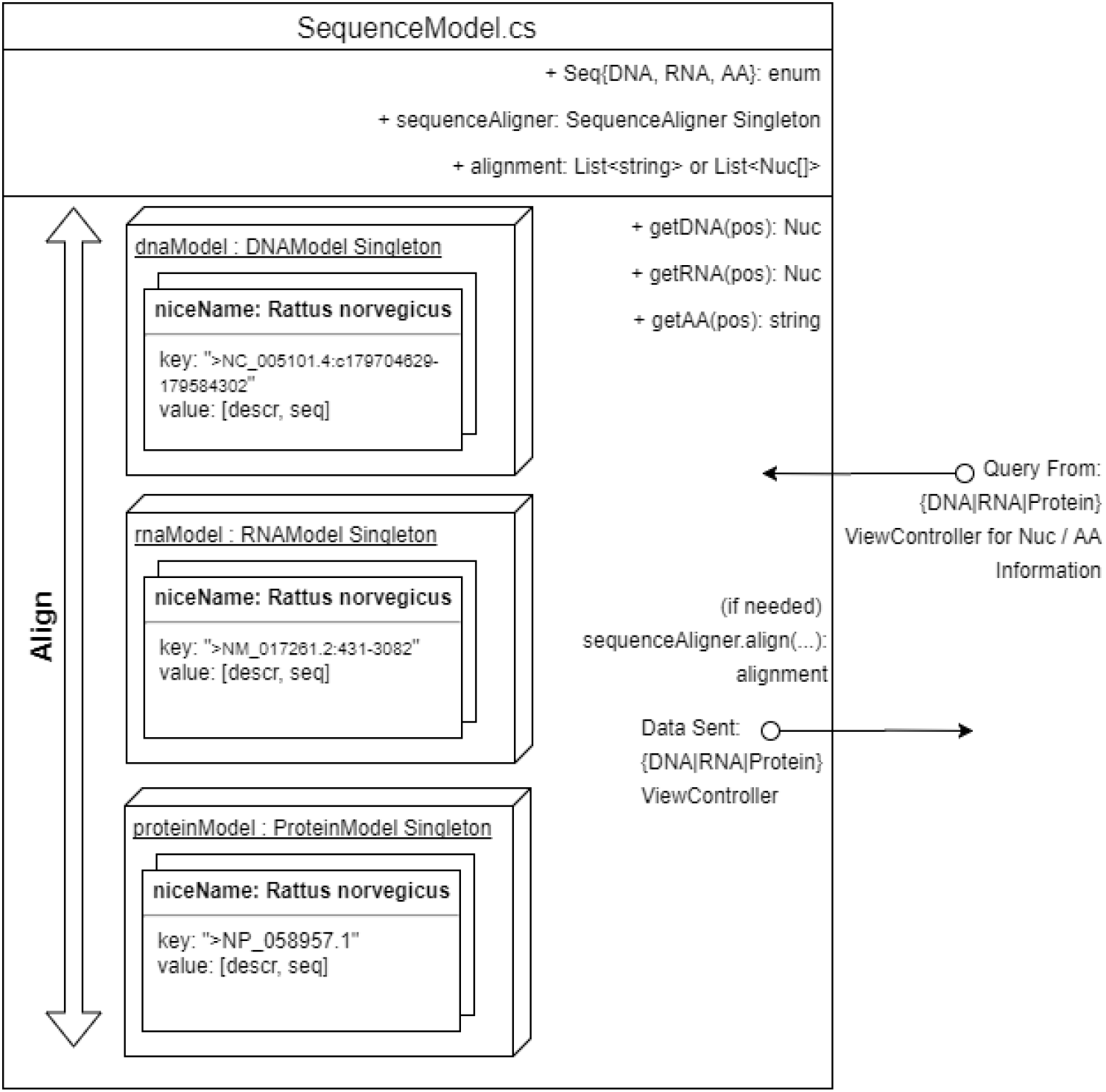
Implementation of the sequence model *SequenceModel*.The model keeps references to DNAModel, RNAModel, ProteinSeqModel singletons, and a reference to a SequenceAligner instance that can implement the Needleman-Wunsch algorithm for global sequence alignment.

#### (E) UI

Hover UI Kit is a free software package available for download from github. It is governed by a GPLv3 license which authorizes free use for open source projects. Once Hover UI libraries are imported into the project directory, an empty GameObject is created, and a Hover UI creation script component is attached. Running the script directly in Unity editor mode results in a static menu set, if given appropriate parameters. The static menu instance is directed to find and attach itself to an instance of Leap Motion hands at runtime (Supplementary Figure S3). The left hand transform acts as the parent to the UI menu while the right hand acts as a pointer. The complete menu hierarchy as implemented in BioVR is listed in Supplementary Table S3.

#### (F) Plane Geometry and UV Coordinates

We use plane geometry and UV coordinates to map nucleotide sequences onto UI elements. A Unity plane geometry is a flat surface as defined by variables contained in its mesh instance. Every mesh instance has an array of Vector2 UV coordinates (Figure 2) which define UV the mapping of a two-dimensional texture image onto the projected surface of any valid geometry. In the case of plane geometry, UV coordinates map perfectly to the length and width of the plane such that in the default case, the U coordinate ranges from (0, 1) and spans the plane’s length, whereas the V coordinate ranges from (0, 1) and spans the plane’s width.

To render nucleotide sequences onto textures, we take advantage of two properties. First, geometries have the unique property of being allowed to partially map to UV coordinates. In other words, mesh geometries do not need to span the entire UV range. Second, Unity allows the procedural editing of textures via *Texture2d.SetPixel*(int x, int y, Color color) where x, y refers to a texture coordinate. Note that a 256 x 256 texture will map to a 1×1 UV square such that (0,0) => (0,0) and (256, 256) => (1,1). Thus, each nucleotide within a sequence can be represented by their traditional colors (Supplementary Table S4) and associated with a specific (u, v) coordinate.

Since geometries don’t need to map to the entire UV space, the *MeshRenderer* component of the plane geometry mesh will then only render portions of the nucleotide sequence at a time (Supplementary Figure S4). The range of the nucleotide sequence to be rendered can be adjusted via scrolling.

### Test datasets

We used the sequences and structures of glutamate ionotropic receptor AMPA type subunit 2 (Gria2) as the test dataset of BioVR as tertiary structure of this protein is available to accompany the genetic sequence. This subunit is one of the four Gria subunits (Gria1-4) involving in the assembly of cation channels (glutamate receptors) activated by alpha-amino-3-hydroxy-5- methyl-4-isoxazole propionate (AMPA). Glutamate receptors play a vital role in mediating excitatory synaptic transmission in mammalian brain and are activated in a variety of normal neurophysiologic processes. Prior studies have identified glutamate binding and closing mechanisms using PDB: 1FTO [25] with the caveat that 1FTO only captures the ligand binding core of the protein. Of the five structures submitted to PDB, 5L1B shows the Gria2 structure in the apo state; therefore 5L1B was selected for use in this project due to its symmetry and unbound nature. Source files containing the Gria2 structure (5L1B) were downloaded from Protein Data Bank (PDB).

Both sequence and structure data of Gria2 from *Rattus norvegicus* are preloaded into the StreamingAssets folder of the BioVR project. BioVR builds have access to a compressed version of the StreamingAssets folder at runtime. During the User Interface Case Study, we plan to provide the equivalent of “built-in” access to these documents for subjects randomized to the Traditional Computer group. A Chrome browser will be open with tabs corresponding to the NCBI page for these documents.

## Results

### Gria2 in BioVR: a test case

BioVR allows scientists to view, at the same time, DNA, RNA and protein (called AA in UI) data. In our test case we visualize the *Gria2* gene in a VR-assisted visualization platform. It features a simple user interface for viewing genomic information which utilizes Leap Motion tracking to turn fingers into data manipulators. To the right hand is attached an instance of Hover UI. The user can select to view DNA, RNA or protein sequences (Figures 5–6).

**Figure 5.**
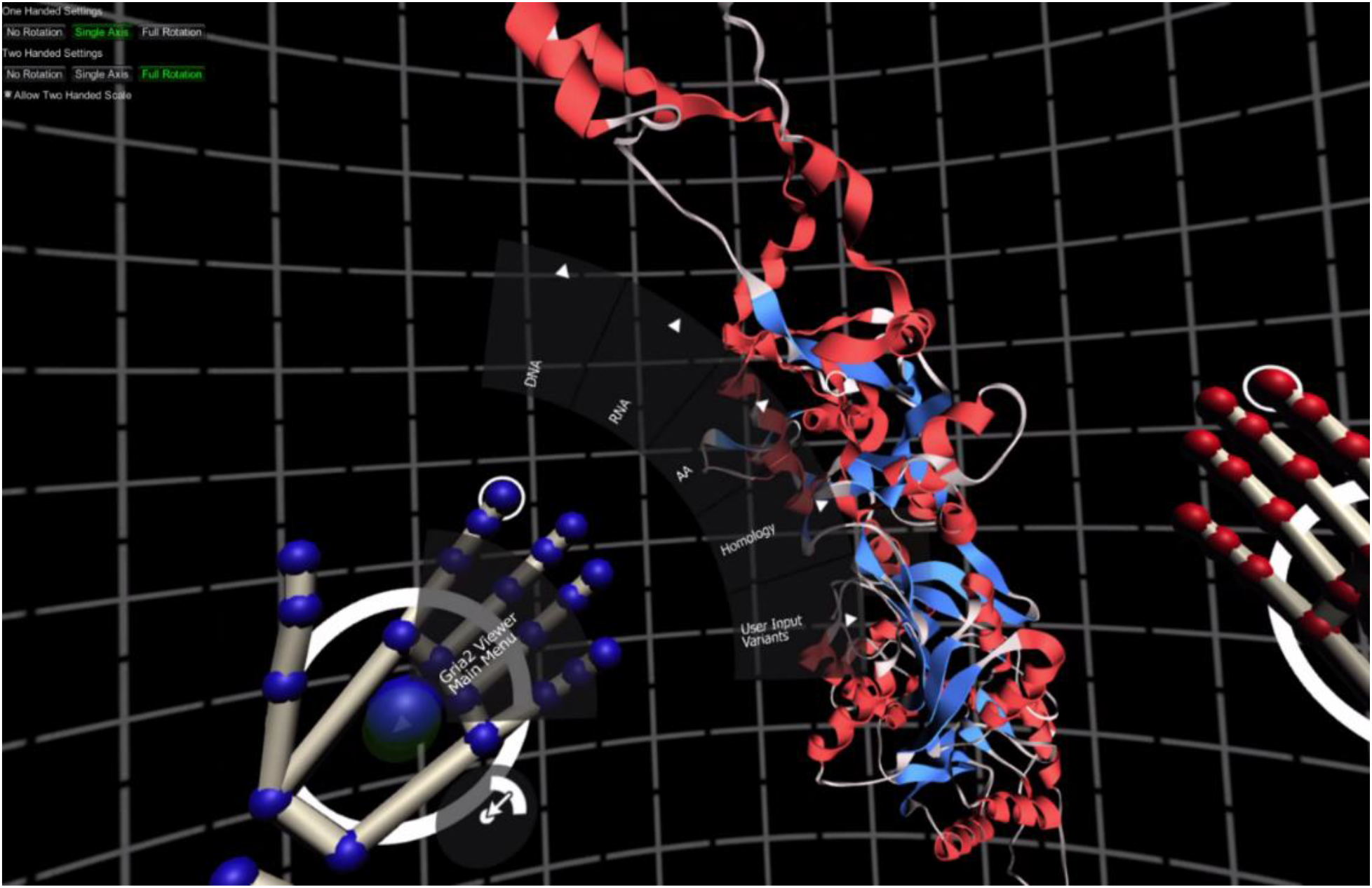
Navigation within BioVR. BioVR uses a menu set anchored to the user’s left hand. The menu set is an instance of Hover UI, an open source project found at https://github.com/aestheticinteractive/Hover-UI-Kit.

**Figure 6.**
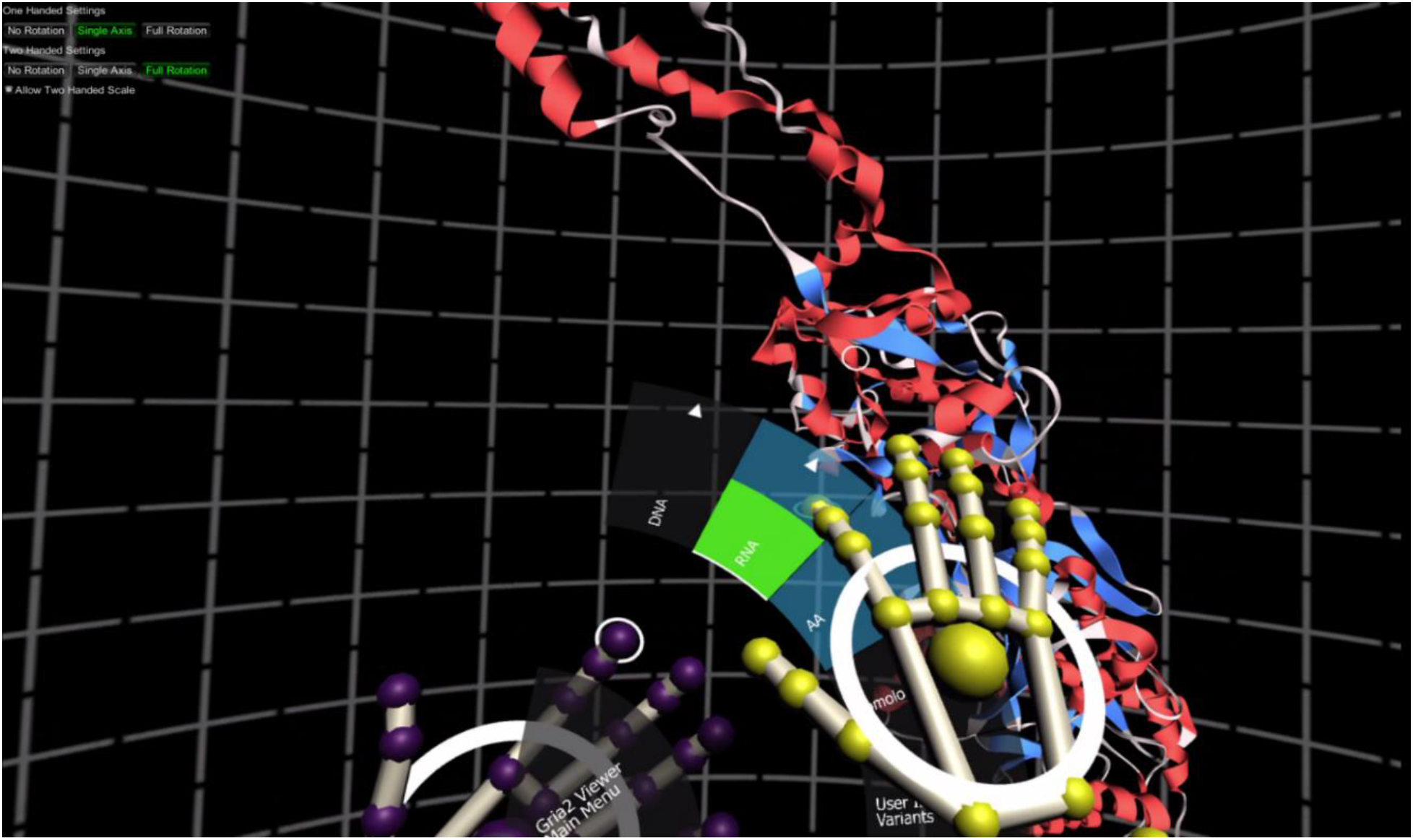
Hover UI used in conjunction with Leap Motion. The menu is anchored to the user’s left hand. The index finger on the right hand acts as a cursor: it generates a button pressed event for a particular button when it hovers over that button within a set amount of time. The specific timing varies and can be set per button.

### Nucleotide Sequences and UV Coordinates

The ideal representation of large (~100 Kb) nucleotide sequences is an open problem in both VR as well as web interfaces. We created a plane geometry with customized UV coordinates that allow for the representation of nucleotide sequences of up to 100 Kb (Supplementary Figure S4). Using the *BuildTexture()* method in {DNA | RNA}PanelController, the appropriate FASTA file is accessed and its nucleotide sequence is processed such that each texture coordinate of the DNA or RNA plane takes on a color that represents a specific nucleotide in the FASTA sequence. The protein sequence in one letter code can be overlaid on top of the mRNA panel (Figure 7). If the user selects “AA>Show on Model” and goes to “RNA>Show”, BioVR gives additional context. Specifically, it displays the 3D structure together with the mRNA sequence of Gria2; the position of selected nucleotide and its corresponding residue on the 3D structure are shown (Figure 7).

**Figure 7.**
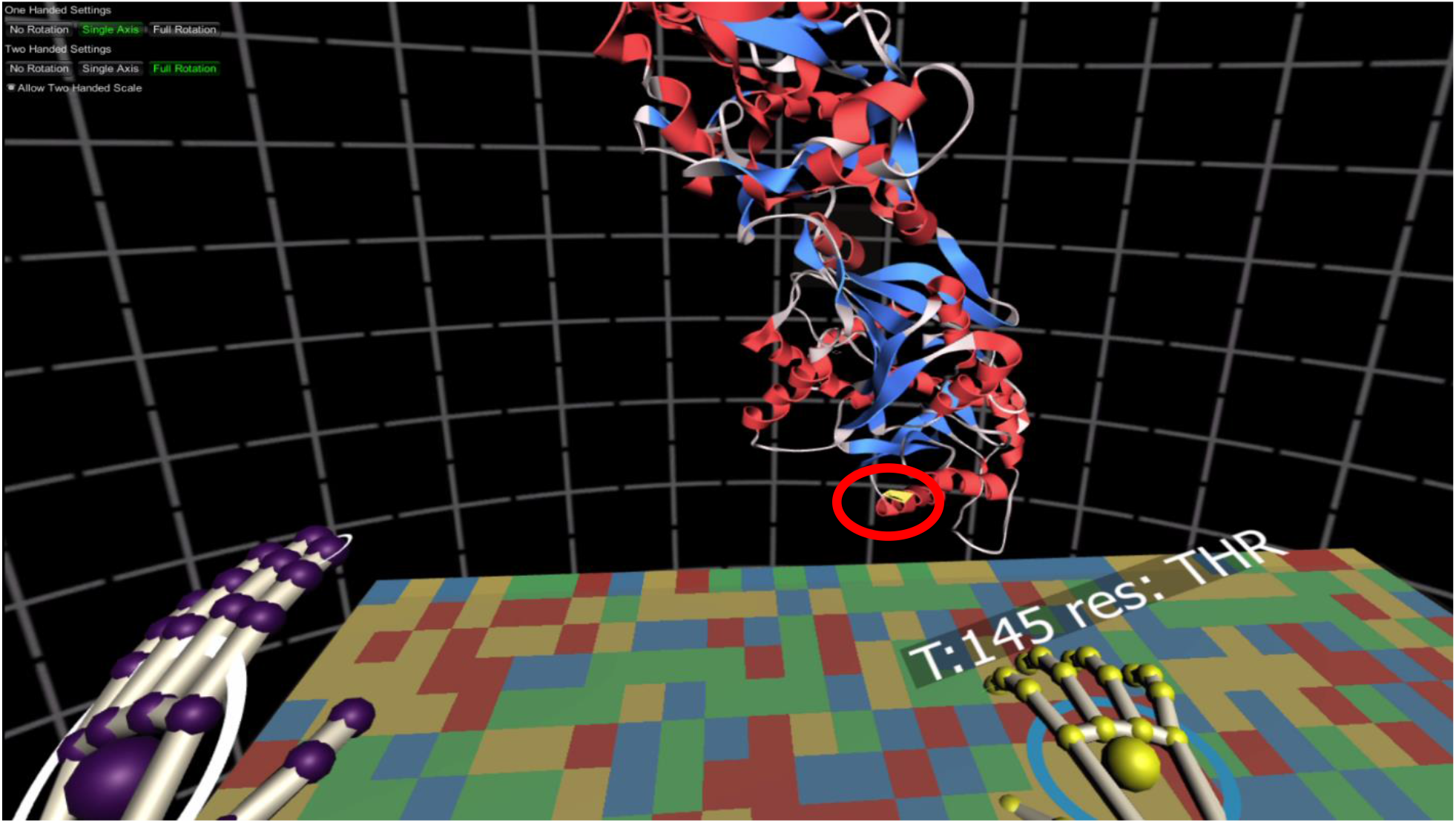
*Rattus norvegicus Gria2* RNA sequence and protein structure shown in BioVR. The mRNA sequence of rat *Gria2* (NM_001083811) was loaded in the RNA Panel. Above the right index finger is a GUI which displays context-sensitive data depending on where the user places his or her index finger along the sequence. The residue corresponding to the mRNA nucleotide is also highlighted in the 3D structure (PDB ID: 5L1B) via a yellow outline shader (denoted by red circle). Note that the red circle was added for the illustration purpose and does not appear in the actual program.

## Discussion

UCSC’s Genome Browser [1], Ensembl Project [2], and Integrative Genomics Viewer (IGV) [3] represent the main genome browsers available to research scientists. In proteomics, UCSF Chimera [26], PyMol [27], and JMol [28] are available to view protein structures. Currently there is a lack of tools to integrate both sequence and structure information. Users have to switch different tools back and forth to gather information they need. This makes comparisons among the various software offerings and their respective UI paradigms difficult to achieve.

In the fields of informatics and biology, new algorithms or experimental procedures gain widespread acclaim if they lead to orders of magnitude increase in research productivity.

Through everyday use of computers for both casual and professional purposes, we intuit that user interfaces have a tangible influence on efficiency. Milestones within the technology industry are often marked by major user interface innovations. For example, from the mouse in Windows to touchscreen interfaces, UI has always had profound an impact on user growth and consumer adoption.

In our view, efficiency gains in genomic analysis could occur during the data visualization stage. Previous studies have shown that researchers tend to understand data more quickly in VR environments than in conventional 2D and 3D desktop visualization [6-8]. Therefore, in this study, we have developed an easy-to-use, VR-assisted platform, BioVR, for visualization of DNA/RNA sequences and protein structures in aggregate. We have shown that DNA/RNA sequences and protein structures can be quickly linked in BioVR. This platform can be extended to view genomic, transcriptomic and proteomic data. This innovation in UI may enhance researchers’ ability to deduct meaningful biological information from more complex datasets, including Hi-C and epigenomic data.

## Conclusions

Virtual reality is a ground-breaking medium with major advantages over traditional visualization for biological datasets. Its potential remains largely unexplored. Here, we develop an easy-to-use, VR-assisted platform, BioVR, and show that DNA/RNA sequences and protein structures can be viewed in aggregate, leading to a novel workflow for researchers.

Our work can be extended to view MPS data to identify SNPs and genomic regions of interest. Whole genome representation can be rendered at low resolution until users decide to investigate a gene locus of interest, upon which the VR application may zoom in to the region at higher resolution. Interactions among promoters, enhancers, and silencers at the loci can be shown depending on chromatin context. Geometric topologies within chromatin regions will be made obvious to the user. Finally, animated simulations can be made to help users visualize temporal datasets.

## Availability and requirements

**Project name**: BioVR

**Project home page**: https://github.com/imyjimmy/gria2-viewer

**Operating system**: Windows 10

**Programming languages**: C#

**Other requirements**: Unity 5.4.2f, Leap Motion, Oculus SDK 1.11.0, Hover UI Kit 2.0.0 beta

**License**: GNU General Public License

## List of abbreviations

UI: user interface
MPS: massively parallel sequencing
*Gria2*: Glutamate Ionotropic Receptor AMPA type subunit 2
HMD: head-mounted display
MVC: Model-view-controller
SoC: separation of concerns

## Acknowledge

Not applicable

## Declarations

### Availability of data and materials

The latest version of the BioVR software is freely available at https://github.com/imyjimmy/gria2-viewer including source code. The Gria2 DNA and RNA sequences are available in NCBI (NM_001083811, NC_005101), and its protein structure is available in PDB (ID: 5L1B).

### Authors’ contributions

JFZ developed the presented software. JFZ and FC drafted the manuscript. ARP provided sequence data. JFZ, ARP, GRS, PAC, JMG and FC edited the manuscript. All authors read and approved the final manuscript.

### Ethics approval and consent to participate

Not applicable

### Consent for publication

Not applicable

